# Machine Learning-Based Identification of Survival-Associated CpG Biomarkers in Pancreatic Ductal Adenocarcinoma

**DOI:** 10.1101/2025.03.29.646090

**Authors:** Yu Zhang, Yining Zhao, Bo Aber Zhang

## Abstract

Pancreatic ductal adenocarcinoma (PDAC) is an exceptionally aggressive cancer with a 5-year survival rate of less than 10%, driven by late-stage diagnosis, limited treatment options, and a lack of reliable biomarkers for early detection and prognosis. In this study, we integrated DNA methylation data from TCGA and ICGC cohorts, categorizing samples based on survival time, and identified 684 differentially methylated CpG sites, along with 224 CpG biomarkers significantly associated with patient survival through statistical and machine learning-based analyses. We developed a random forest model to predict patient survival, achieving 85.2% accuracy for short-survival patients and 70.0% for long-survival patients in the validation set. External dataset validation further confirmed the model’s robustness and accuracy. *De novo* motif analysis of genomic regions surrounding the 224 CpG biomarkers identified *TWIST1* and *FOXA2* as key transcriptional regulators enriched in survival-associated CpG sites, linking their activity to patient survival outcomes. Collectively, our findings highlight valuable epigenetic biomarkers and provide a predictive model to assess PDAC risk levels post-surgery, offering the potential for improved patient stratification and personalized therapeutic strategies.

## Introduction

Pancreatic cancer, particularly pancreatic ductal adenocarcinoma (PDAC), is among the most aggressive and lethal malignancies, characterized by late-stage diagnosis and limited treatment options. Despite advancements in research, the survival rate for PDAC remains dismal, with a 5-year survival rate below 10%^1,2^, and 23.3% of patients experiencing relapse within 12 months post-surgery^3^. To improve patient outcomes, identifying reliable biomarkers for early detection, prognosis, and personalized treatment has been a significant focus in recent years.

Biomarkers derived from genetic, epigenetic, transcriptomic, and proteomic indicators are critical for improving diagnostic precision and predicting patient outcomes. However, the heterogeneity of pancreatic cancer, both genetically and epigenetically, complicates the discovery of robust, generalizable biomarkers. Recent advances in machine learning (ML) have provided powerful tools to analyze high-dimensional data and identify meaningful patterns in biomarker research. ML approaches have shown high accuracy in pancreatic lesion classification and early detection of pancreatic cancer. Deep learning has been applied to imaging analyses, including cross-sectional imaging^4^ and non-contrast computed tomography^5^. Similarly, ML has been instrumental in identifying circulating protein biomarkers, such as CA19-9, and inflammatory proteins to distinguish PDAC patients from non-PDAC individuals^6^. Predictive biomarkers like cytokines (eotaxin-3, *IL-10*) and immune cell dynamics (CD4+ and CD8+ lymphocytes) have been linked to survival and treatment efficacy^7^. Additionally, exosomal microRNAs (miRNAs) have emerged as promising diagnostic tools for early detection and therapeutic targets^8^. Genetic mutations, such as *SMAD4* deficiency, have further provided insights into therapeutic resistance and prognostic potential^9^.

DNA methylation biomarkers have garnered attention as promising non-invasive tools for early detection and monitoring of PDAC progression^10-13^. Aberrant methylation in circulating tumor DNA (ctDNA) of tumor suppressor genes has demonstrated potential for early-stage detection, although specificity remains a challenge when used independently^14-16^. A four-gene DNA methylation signature (*CCNT1, ITGB3, SDS, HMOX2*) has shown superior accuracy in stratifying high-risk and low-risk patients for survival prediction^17^. Machine learning models, such as random forest, have achieved high accuracy in detecting non-ductal pancreatic neoplasms using DNA methylation patterns^18^. Methylation-associated alterations in pathways regulating oxidative stress, DNA repair, and cell cycle have been linked to distinct PDAC subtypes, improving our understanding of tumor biology and prognosis^19,20^. Multi-omics analyses of cfDNA methylation have further identified novel biomarkers with high potential for early PDAC detection and diagnostic precision^21^. These findings underscore the transformative potential of biomarkers in PDAC diagnosis and treatment. By combining machine learning techniques with multi-omics analyses, researchers are accelerating the discovery and validation of biomarkers, paving the way for more effective clinical applications and personalized medicine in PDAC.

In this study, we analyzed DNA methylation data from TCGA and ICGC cohorts to explore survival-associated patterns in pancreatic cancer. By grouping samples based on survival durations, we identified 684 CpG sites with differential methylation and pinpointed 224 CpG biomarkers closely tied to patient outcomes using advanced statistical methods and machine learning approaches. We further developed a random forest model to predict the survival status of PDAC patients, demonstrating 85.2% accuracy for short-survival patients and 70.0% for long-survival patients in the validation dataset. The model’s performance was further validated using external datasets, confirming its reliability and generalizability. Further investigation into the genomic regions surrounding these 224 CpG biomarkers revealed significant enrichment for the transcription factors *TWIST1* and *FOXA2*, linking their regulatory influence to survival-related DNA methylation changes. Our research not only identifies promising biomarkers for pancreatic cancer but also provides a predictive framework to assess post-surgical risk, paving the way for enhanced patient stratification and more tailored therapeutic interventions.

## Result

### DNA methylation profiles that distinguish long-survival from short-survival PDAC patients

To better understand the DNA methylation profiles that differentiate long-survival (LS) from short-survival (SS) pancreatic ductal adenocarcinoma (PDAC) patients, we integrated DNA methylation data (Illumina Infinium 450K) from TCGA PDAC samples with Australian PDAC samples from the ICGC. In total, 281 samples (Supplementary table 1) were included and categorized into three survival groups: 118 Short Survivors (SS, <12 months), 110 Mid Survivors (MS, 12-36 months), and 53 Long Survivors (LS, >36 months), excluding patients still alive in the SS and MS groups (Fig. 1a). Principal component analysis revealed that global DNA methylation profiles of PDAC patients were largely independent of survival time (Supplementary Fig. S1a).

**Figure 1.**
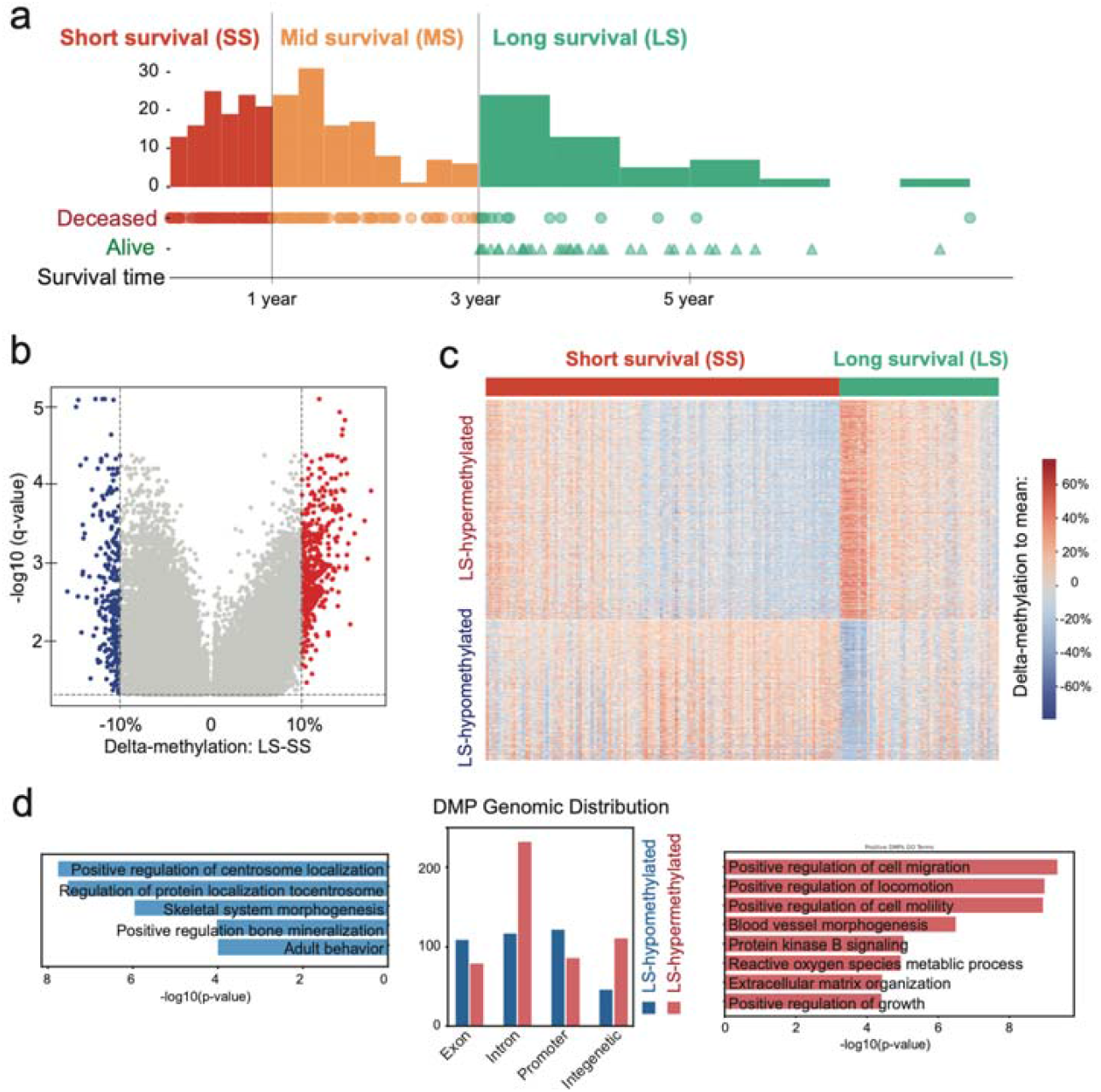
Differentially methylated CpG positions (DMPs) can distinguish long-survival patients from short-survival patients. **a)**. Distribution of survival time and deceased status in TCGA and ICGA PDAC patients. **b)**. Volcano plot of DMPs between long-survival (LS) patients and short-survival (SS) patients. **c)**. DNA methylation heatmap of 668 DMPs in all long- and short-survival patients. d). Gene Ontology biological process enriched in LS-hypomethylated DMPs tagged genes (left) and LS-hypermethylated DMPs tagged genes (right), with distinct genomics distribution (middle).

We identified 688 differentially methylated CpG (DMC) sites between the SS and LS groups, including 420 hypermethylated and 268 hypomethylated DMCs in the LS group compared to the SS group (Fig. 1b). Despite some heterogeneity, distinct DNA methylation patterns were evident among the identified DMCs in the patients (Fig. 1c). Distance analysis of DMCs from the nearest genes showed that 58% were located within 5KB of transcription start sites, while only 207 DMCs were directly within gene promoter regions (Fig. 1d, Supplementary Fig. S1b). Gene ontology analysis indicated that genes near LS-hypermethylated DMCs were primarily associated with centrosome regulation and function. Additionally, LS-hypermethylated DMCs were linked to processes such as reduced cell migration and mobility, blood vessel morphogenesis, and the suppression of cancer cell growth (Fig. 1d). These findings suggest that genes promoting cancer cell growth and migration are less active in long-survival PDAC patients compared to short-survival patients.

### Identification of the CpG biomarkers associated with the survival time of PDAC patients

To identify CpG biomarkers associated with the survival time of PDAC patients, we employed two complementary strategies: correlation regression and random forest analysis, to pinpoint the most relevant CpG sites (Fig. 2a). Using DNA methylation data from TCGA and ICGC for PDAC patients, we combined the LS and SS groups and randomly split the dataset into a training and validation set (80%) and a test set (20%), resulting in 134 samples in the training and validation set and 37 samples in the test set. Additionally, we incorporated the MS group (90 patients) into the training and validation set and calculated the Pearson correlation coefficient between DNA methylation levels and patient survival times for each CpG site (Fig. 2b). The 3,000 CpG sites with the most significant correlation coefficients were selected as the candidate pool for further analysis.

**Figure 2.**
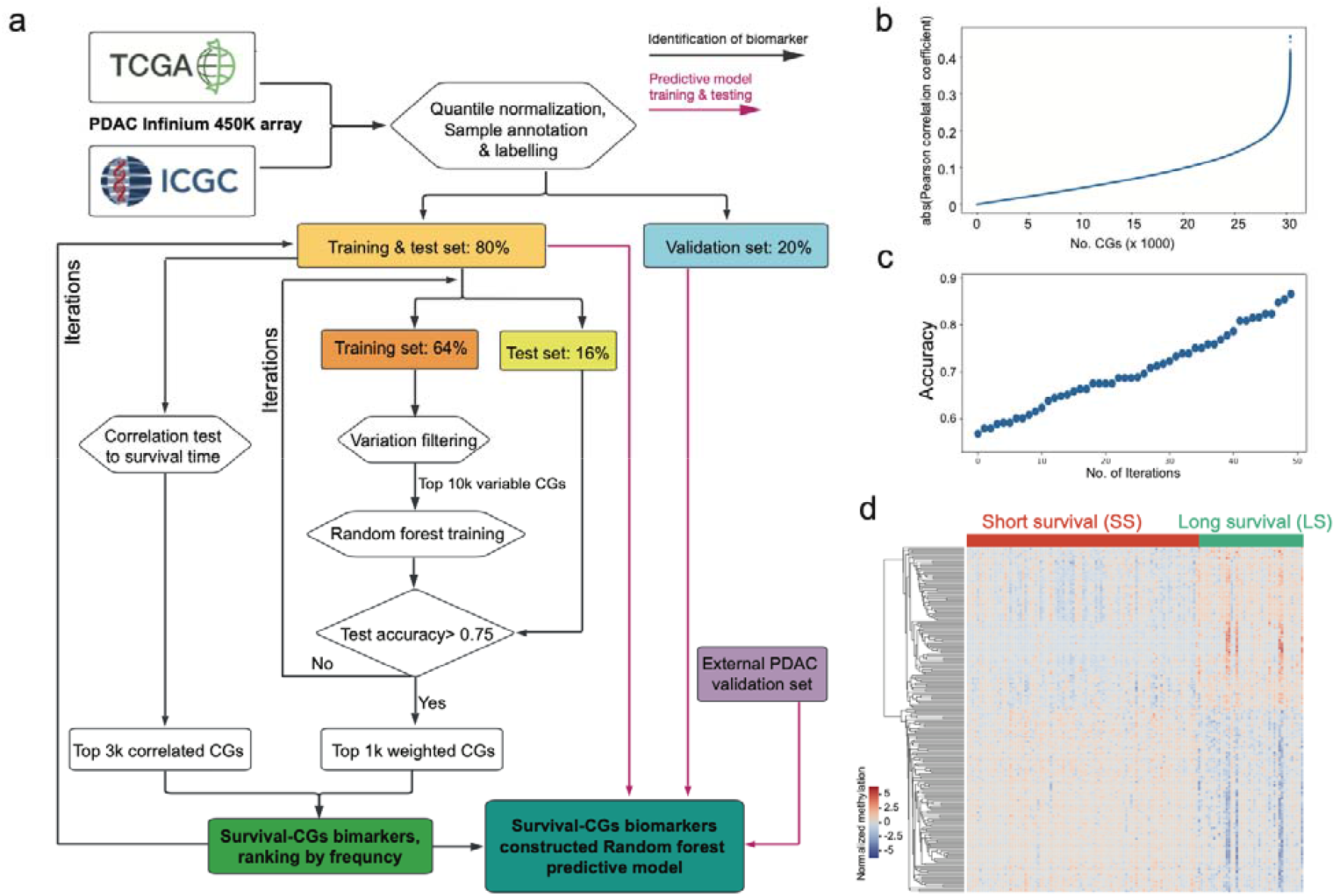
Identification of PDAC patient survival-associated CpG biomarker. **a)**. Flowchart of Identification of PDAC patient survival-associated CpG biomarkers through machine-learning approaches. **b)**. absolute Pearson correlation coefficient of all CpG sites against the survival times in TCGA and ICGA dataset. **c)**. Ranked test accuracy in 50 times Random Forest training (feature selection). **d)**. DNA methylation heatmap of 224 survival-associated CpG biomarkers in all long- and short-survival patients.

We further refined the selection of CpG biomarkers by applying random forest decision models for feature selection. The combined training and test set, which included 80% of SS and LS patients, was further randomly divided into a training set and a test set at an 80/20 ratio, representing 64% and 16% of the total SS and LS patients, respectively. Using the training set, the top 10,000 most variable CpG sites were identified and used to train a random forest classification model. The performance of the trained model was evaluated on the test set, and the top 1,000 CpG sites with the highest variable importance were retained, provided the prediction accuracy exceeded 75% (Fig. 2c).

We then intersected the 3,000 CpG candidates with the most significant Pearson correlation coefficients with the 1,000 CpG sites identified as having the greatest variable importance in the random forest model. Through 100 successful iterations of this process, we identified 224 CpG sites with the highest appearance frequency as robust biomarkers associated with PDAC survival (Supplementary table 2). Notably, nearly half of these biomarkers exhibited hypomethylation in the long-survival group, highlighting their potential role in influencing favorable survival outcomes (Fig. 2d). These results emphasize the utility of combining statistical correlation and machine learning approaches to identify meaningful survival-associated biomarkers.

### Methylation profiles predict the survival of PDAC patients

Using the identified 224 CpG biomarkers associated with the survival of PDAC (pancreatic ductal adenocarcinoma) patients, we trained a random forest model utilizing 80% of the LS (long-survival) and SS (short-survival) patient cohorts with 4-fold cross-validation (Fig. 2a). The model consists of 500 decision trees, each incorporating different combinations of selected CpG biomarkers, as illustrated in one example tree (Fig. 3a). By analyzing the DNA methylation levels in the top three layers of the decision tree, we observed distinct patterns for each CpG site, corresponding to short-survival, mid-survival, and long-survival patient groups (Fig. 3b). This demonstrates the discriminatory power of DNA methylation levels in stratifying survival outcomes.

**Figure 3.**
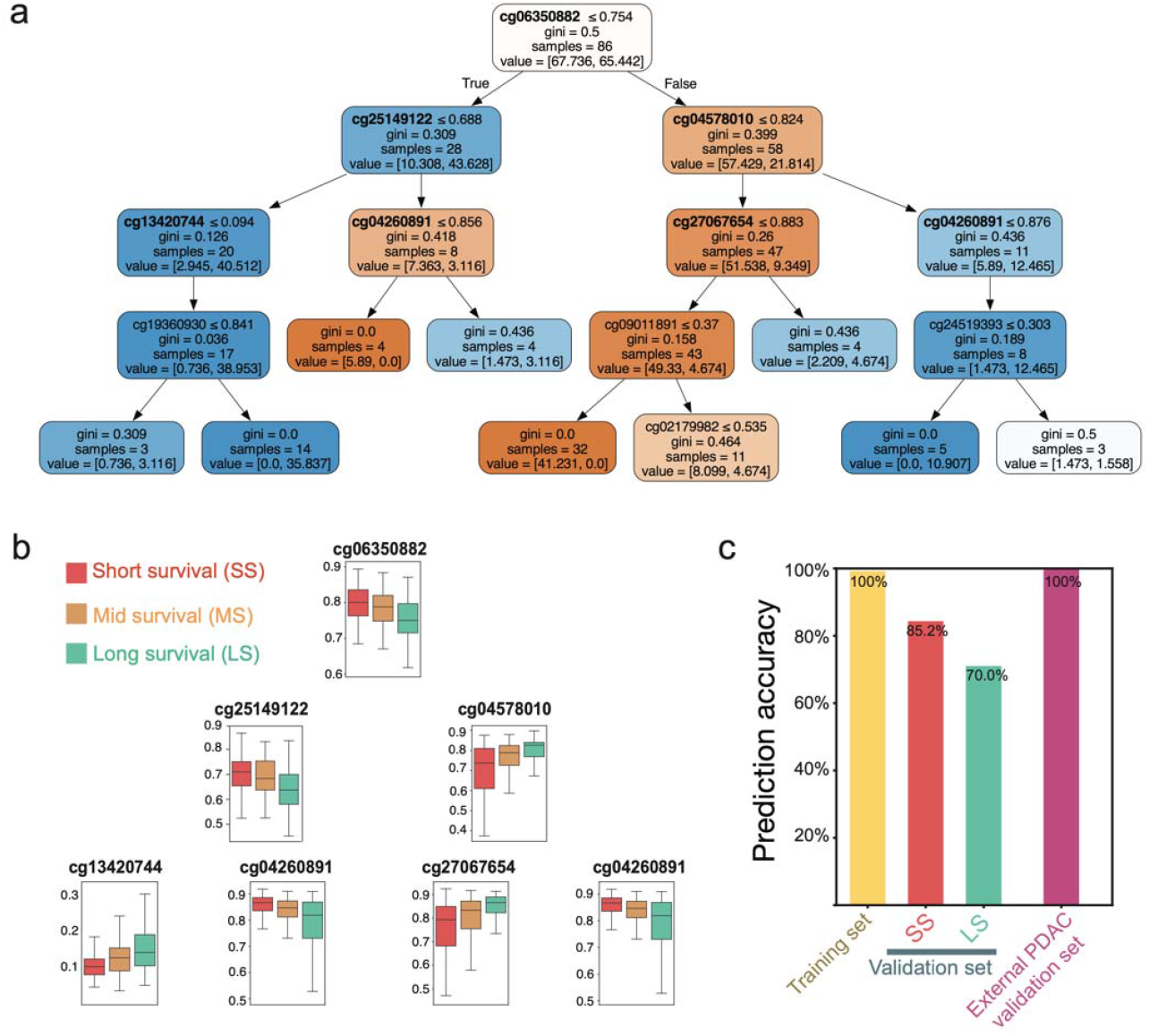
Random Forest predictive model based on 224 survival-associated CpG biomarkers. **a)**. 1^st^ decision tree of select Random Forest predictive model. **b)**. DNA methylation level of CpGs used in the 1^st^ decision tree in short-, middle-, and long-survival PDAC patients. **c)**. Prediction accuracy of selected Random Forest predictive model on training and validation PDAC dataset.

Genomic distribution analysis revealed that 21% of the 224 CpG biomarkers are located within distal regulatory elements, while the remaining are near gene promoters (Supplementary Fig. S1c, Supplementary table 3).

Notably, cg06350882, situated in the top layer of the decision tree, is strongly associated with survival outcomes. Patients with lower DNA methylation levels at this CpG site exhibited longer survival times. This CpG site is located in a highly conserved intergenic enhancer region, far from any known protein-coding genes, suggesting a potential regulatory role (Supplementary Fig. S2).

We further evaluated the performance of the trained random forest-based predictive model (supplementary file 1) using the validation set, which comprised the remaining 20% of TCGA and ICGC samples (Fig. 3c). During the training phase, the model achieved 100% accuracy in distinguishing short-survival and long-survival patients within the training dataset, demonstrating its ability to effectively learn survival-associated DNA methylation patterns. In the validation dataset, the model correctly identified 23 out of 27 short-survival PDAC patients (accuracy: 85.2%) and 7 out of 10 long-survival PDAC patients (accuracy: 70.0%). These results indicate the model’s high accuracy in recognizing DNA methylation patterns linked to survival outcomes. To further assess its robustness, we applied the trained model to an external independent PDAC DNA methylation dataset from a previous study^22^, which included survival data for each patient. In this external dataset, the model achieved 100% accuracy for short-survival patients (n=5) (Fig. 3c). However, for very limited long-survival patients (n=2), the predictive model failed (Supplementary table 3). These findings suggest that while the model is highly sensitive and reliable in identifying short-survival samples, it exhibits reduced sensitivity to DNA methylation patterns associated with long-survival patients.

We compared the 224 survival-associated CpG biomarkers with differentially methylated CpG sites (DMCs) identified between the long-survival (LS) and short-survival (SS) groups. Only 60 CpG sites were shared between the two groups (Fig. 4a), suggesting that the DNA methylation signatures associated with survival and group-specific differences represent distinct epigenetic characteristics in pancreatic cancers. This distinction highlights the complexity of DNA methylation patterns in influencing survival outcomes. Gene ontology (GO) analysis of the genes associated with the 224 survival-related CpG biomarkers revealed significant enrichment in biological processes such as cell migration, actin filament organization, and DNA replication (Fig. 4b). These pathways are closely linked to the aggressiveness of cancer cells, underscoring the relevance of these genes in pancreatic cancer progression and metastasis.

**Figure 4.**
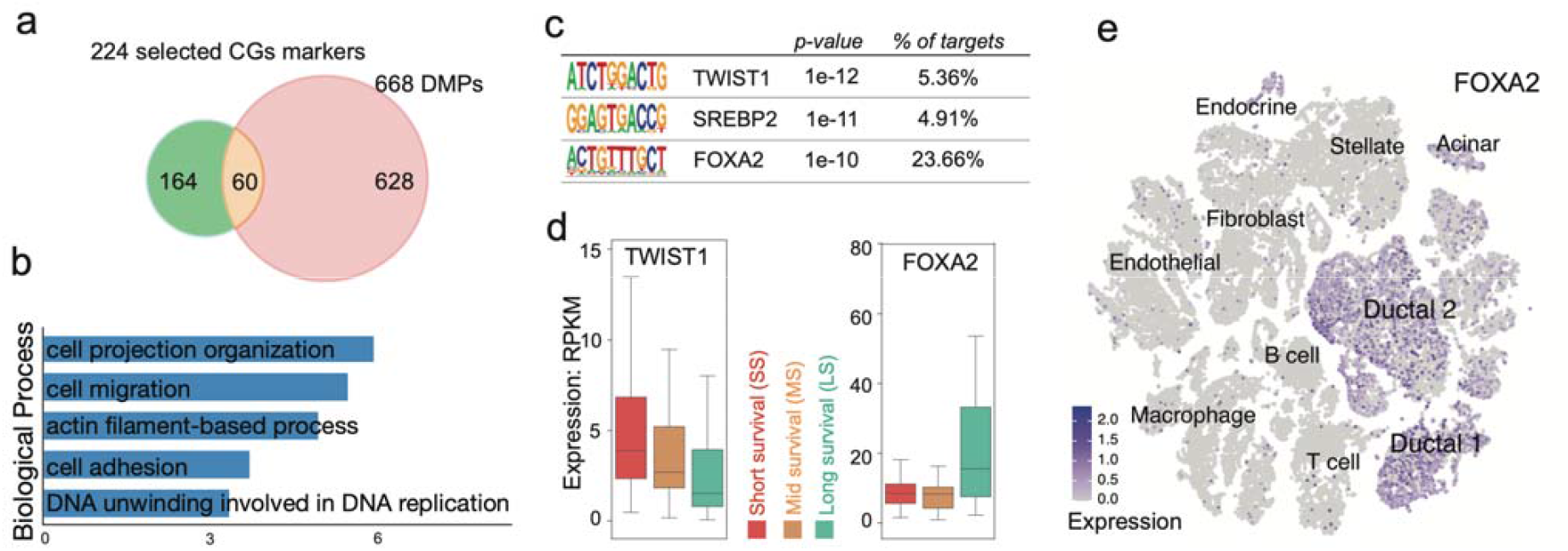
Survival-associated CpG biomarkers connect PDAC patients’ survival to key transcription factors (TFs). **a)**. Shared CpG sites between survival-associated CpG biomarkers and DMPs. **b)**. Gene Ontology biological process enriched in Survival-associated CpG biomarkers tagged genes. **c)**. TFs enriched in the regions containing Survival-associated CpG biomarkers. **d)**. Expression of *TWIST1* and *FOXA2* in short-, middle-, and long-survival PDAC patients. **e)**. *FOXA2* expression in different cancer cell types of 24 PDAC tumors.

DNA methylation is often influenced by transcription factor (TF) binding at genomic regions. To explore this, we performed *de novo* motif analysis on 200bp regions surrounding the 224 CpG biomarkers and identified three highly enriched transcription factors: *TWIST1, SREBP2*, and *FOXA2* (Fig. 4c). Further investigation into the expression levels of these TFs in matched TCGA patient samples revealed significant correlations between *TWIST1* and *FOXA2* expression and patient survival time (Fig. 4d). Elevated expression of *TWIST1* was observed in short-survival patients and was negatively correlated with survival time (r=-0.19). As a well-established oncogene, *TWIST1* promotes cancer cell invasion and dissemination through epithelial-mesenchymal transition (EMT)^23,24^. It is also associated with the stem-like properties of cancer cells, contributing to therapy resistance^25^. Single-cell transcriptome analysis of 24 PDAC patients^26^ further showed that *TWIST1* was predominantly expressed in fibroblast-like cells and pancreatic stellate cells (Supplementary Fig. S3a), suggesting a key role in shaping the tumor microenvironment and supporting cancer progression.

Known as a pioneer transcription factor, *FOXA2* regulates pancreas-specific gene expression during development. Its binding motif was found near 24% of the 224 survival-associated CpG biomarkers, emphasizing its role in metastasis and patient survival. *FOXA2* expression was significantly higher in long-survival PDAC patients (p-value=0.004, two-tailed t-test), and single-cell RNA-seq data revealed its strong ductal-specific expression compared to other pancreatic cancer cell types. This suggests that *FOXA2* may play a protective role in pancreatic cancer by maintaining differentiation and inhibiting aggressive phenotypes. These findings highlight the dual roles of TFs like *TWIST1* and *FOXA2* in modulating the epigenetic landscape and shaping patient outcomes in pancreatic ductal adenocarcinoma (PDAC). TWIST1’s involvement in promoting tumor invasiveness and *FOXA2*’s association with ductal cell identity and metastasis inhibition offer valuable insights into their functional significance and potential as therapeutic targets in PDAC.

Finally, we identified five of the 224 CpG biomarkers located within transposable elements (Supplementary Table 5). These highly repeated DNA sequences, originating from ancestral transposons, used to move and duplicate within the human genome, significantly contribute to the regulatory elements in regulating gene expression^27,28^. One notable example is cg05181795, which is associated with the MER96B transposable element and located upstream of *MDFI* (Supplementary Fig. S4a). *MDFI* (MyoD Family Inhibitor) is overexpressed in multiple cancer types, including colorectal cancer, breast cancer, liver cancer, and lung cancer, suggesting a potential role in tumor progression^29,30^. In PDAC, we noticed an increased DNA methylation level of cg05181795 associated with longer survival time, accompanying with decreased expression level of *MDFI* (Supplementary Fig. S3b). Another key biomarker, cg19360930, is associated with an endogenous retrovirus-like (ERVL) element, LTR102_Mam, located within the 3’ UTR of *WDR35. WDR35* is essential for ciliary formation and elongation, playing a critical role in ciliogenesis and developmental processes. Notably, the LTR102_Mam element harbors multiple transcription factor binding sites and is located 280 bp from a known *FOXA2* binding site (Supplementary Fig. S4b). However, the increased DNA methylation level of cg19360930 along with longer survival time did not correlate with expression of *WDR35* (Supplementary Fig. S3c). These results suggest the necessary future study to understand the transposable element’s involvement in gene regulation during pancreatic carcinogenesis through transcriptional modulation.

## Discussion

DNA methylation alterations are a hallmark of human cancer cells, and have been well studied to be associated with human development^31,32^ and carcinogenesis^33,34^. In this study, we integrated DNA methylation data from the TCGA and ICGC cohorts to investigate survival-associated methylation patterns in pancreatic cancer. Through statistical analysis, we identified 688 CpG sites with differential methylation patterns between long-survival and short-survival patients. Gene Ontology analysis revealed that genes located near LS-hypermethylated DMCs were primarily associated with centrosome regulation and function. Additionally, LS-hypermethylated DMCs were linked to reduced cell migration and mobility, impaired blood vessel morphogenesis, and the suppression of cancer cell proliferation. These findings suggest that the epigenetic landscape of long-survival patients is closely linked to key cancer cell characteristics, indicating that DNA methylation signatures may serve as valuable biomarkers for predicting patient survival, reflecting the influence of distinct cancer cell subtypes on disease progression.

We further applied machine-learning approaches to identify 224 CpG biomarkers closely associated with patient outcomes. Notably, only 60 of these biomarkers overlapped with the 684 differentially methylated positions (DMPs) identified earlier. This result suggests that critical information regarding cancer cell subtypes, disease progression, and patient survival is deeply embedded in DNA methylation patterns. Further investigation of the genomic regions surrounding these 224 CpG biomarkers revealed significant enrichment for the transcription factors *TWIST1* and *FOXA2*, linking their regulatory influence to survival-related DNA methylation changes.

*TWIST1* is a key regulator of epithelial-to-mesenchymal transition (EMT), a process crucial for cancer metastasis and cell migration in multiple cancer types^23,24^. In PDAC, *TWIST1* was recently found to regulate glycolytic metabolism^35^, and was also regulated by *AURKA*^36^ and *TGIF1*^37^. We observed a strong correlation between *TWIST1* expression and patient survival time, suggesting that *TWIST1* expression levels could serve as a potential prognostic biomarker for PDAC (pancreatic ductal adenocarcinoma). *FOXA2* is a master transcription factor that regulates pancreas-specific gene expression during development^38^. Notably, the higher expression levels of *FOXA2* in long-survival patient groups indicate its potential role as a tumor suppressor gene, consistent with previous findings in various cancer types^39-41^, including PDAC^42-45^. Single-cell transcriptome analysis further revealed that *FOXA2* is specifically expressed in ductal cells, highlighting its essential role in maintaining pancreatic ductal cell identity and reinforcing its significance in PDAC tumor biology.

Finally, we further developed a random forest model to fully utilize the distinguishing power of 224 CpG biomarkers associated with patient survival and predict the survival status of PDAC patients. After training and optimizing the model using 80% of the TCGA and ICGC datasets, we evaluated its performance on the remaining 20% of the dataset, which had not been used in training. The model achieved an accuracy of 85.2% for short-survival patients and 70.0% for long-survival patients in this validation dataset. This performance highlights the potential application of our predictive model in assessing and estimating the short-survival risk of PDAC patients immediately after surgery.

To test the robustness of our model across different datasets, we further validated it using an entirely independent dataset from a previous PDAC study^22^. The model demonstrated 100% accuracy in predicting short-survival patients; however, it showed lower predictive accuracy for long-survival patients. When combined with the lower accuracy observed for long-survival predictions in the TCGA/ICGC validation dataset, this result suggests that our training model still has limitations in accurately identifying long-survival cases compared to its highly precise predictions for short-survival patients. This discrepancy may be attributed to the relatively small number of long-survival patients in the training dataset. The imbalanced data distribution likely affects the weight of each decision tree, making the model more sensitive to short-survival predictions. Future efforts will focus on refining the model and incorporating additional data to enhance its sensitivity and overall predictive power for long-survival patients. As more publicly available PDAC methylation datasets with survival time and status become accessible, future iterations of our predictive model will greatly benefit from these expanded resources. Despite these limitations, this predictive approach by using DNA methylation data holds significant promise for real-world applications in PDAC patient care.

## Methods

### Dataset and Pre-processing

DNA methylation data were obtained from the TCGA data portal (https://portal.gdc.cancer.gov) and ICGC data portal(https://dcc.icgc.org), specifically using the Illumina Infinium Human Methylation 450 BeadChip. Following data download, probes with missing beta values in either TCGA or ICGC samples were excluded. Additionally, probes overlapping known dbSNP SNPs with a minor allele frequency (MAF) greater than 0.01 were filtered to minimize the impact of genetic variation on methylation analysis. 323 probes mapping to the Y chromosome were also removed. To correct for technical variation and batch effects arising from the use of distinct datasets, we performed quantile normalization using the *preprocessCore* package in *R*. The resulting dataset comprised 281 tumor samples and 303,899 CpG sites.

### Identification of Differentially Methylated CpGs (DMCs)

To identify differentially methylated CpG sites (DMCs) in pancreatic cancer, we analyzed normalized methylation data from the short-survival and long-survival groups using the *dmpFinder* function in the *minfi* package. For each CpG site, we calculated the q-value and *delta*_(Long – Short)_, representing the mean beta value difference between the two groups. CpG sites were classified as DMCs if they met the cutoff criteria of |*delta*_(Long – Short)_| ≥ 10% and q-value < 0.05. DMCs were further categorized into positive and negative groups (N_positive = 420, N_negative = 268) based on their delta values. Positive *delta*_(Long – Short)_ indicates hypermethylation in the long-survival group, while negative *delta*_(Long – Short)_ indicates hypomethylation in the long-survival group.

### Selection and Identification of Survival-associated CpG biomarkers

To avoid overfitting, following data normalization of PDAC DNA methylation data, we randomly allocated 20% of the total samples to a held-out validation set, which was excluded from further training steps. The remaining samples were further randomly split into a training set (64%) and a testing set (16%) for downstream feature selection, combined with bootstrap sampling iteration strategy. Firstly, we applied correlation-based filtration, to calculate the Pearson correlation between methylation beta values and survival time for each CpG site. The top 3,000 CpG sites with the strongest correlations were selected as the “correlation candidate pool”, and will be used to intersect with candidates selected from the second strategy. The second strategy utilized a between-group variation (BGV) filtering approach followed by Random Forest feature importance. Specifically, BGV was calculated for each CpG site by using the randomly sampled training set (64% of total samples) as follows:

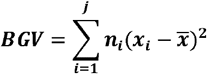

Where ***j*** is the number of groups,*n*_*i*_ is the size of group ***i***, ***x***_***i***_ is the mean of group i, 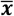 is the grand mean of samples across all groups. In this study, ***j*** *=* 2. To reduce downstream training complexity, the top 10,000 CpGs with the largest BGV were selected as input features for the subsequent Random Forest (RF) training. The RF model was trained by using the training set (64% of total samples), further tested the predictive accuracy by using the testing set (16% of total samples). Once the predictive accuracy achieved ≥ 75%, the Gini importance of CpG features was collected and ranked, and the top 1,000 CpG sites were selected and intersected to “correlation candidate pool”. The CpG features shared between the two strategies were recorded. The random sampling of the training set (64%) and the testing set (16%) were iterated 100 times, followed by the two-strategies-combined machine-learning feature selection after each random sampling. The top 5% of candidates CpGs in the 100 times feature-selection trainings were identified as PDAC survival-associated CpG Biomarkers.

### Predictive Modeling of Survival Time

To predict survival in pancreatic ductal adenocarcinoma (PDAC) patients, we developed a Random Forest classifier. This ensemble method aggregates predictions from multiple decision trees to enhance predictive accuracy and robustness. 224 survival-associated CpG sites, identified through feature selection (as described previously), were used as input features for the model. For model training and evaluation, the previously partitioned training (64%) and testing (16%) sets were combined. We employed a grid search approach with 3-fold cross-validation on the combined dataset to optimize model hyperparameters. The hyperparameters explored included the number of trees (n_estimators), minimum samples split (min_samples_split), and minimum samples leaf (min_samples_leaf). The final model was selected based on the hyperparameter combination that yielded the best performance, as assessed by highest training accuracy.

The held-out validation set (20% of total samples) and one external PDAC DNA methylation dataset^22^ were used to evaluate the accuracy and robustness of the trained RF predictive model. DNA methylation data of Illumina 450K array was downloaded from GEO (GSE165687), and data cleaning and normalization were performed as described above. Only the patients followed the short or long survival standards defined above were used for validation. The DNA methylation beta-values of the matched 224 survival-associated CpG biomarkers were input into the trained RF predictive model.

### Gene Ontology (GO) Analysis and motif analysis

GO enrichment analysis of identified DMCs and survival-associated CpG biomarkers was performed by using ToppGene Suite^46^. The genes associated with DMCs and survival-associated CpG biomarkers were identified by using the annotation table of Illumina Infinium 450K array. Significant enriched biological process terms were selected with cutoffs of FDR<0.05.

The transcription factors binding motifs (TFBS) enriched in the regions around survival-associated CpG biomarkers were analyzed by using findMotifsGenome.pl (-size given) of HOMER^47^ software (v4.11.1). The 200bp region around each CpG biomarker in the human genome was used as potential TF binding regions. The significantly enriched known binding motifs in regions were identified with p-value cutoff <1e−10.

### TCGA RNA-seq data processing

RNA-Seq data from patients with Pancreatic Ductal Adenocarcinoma (PDAC) were retrieved from The Cancer Genome Atlas (TCGA) data portal, accessible at https://portal.gdc.cancer.gov. To investigate the relationship between gene expression and patient prognosis, we applied the same classification criteria as described previously [cite previous study or section]. Specifically, patients were stratified into three groups based on their overall survival (OS) time: short survival (SS) [survival time< 1 year]; medium survival (MS) [1 year ≤ survival time < 3 years]; and long survival (LS) [survival ≥ 3 years]. Gene expression levels were quantified using the Fragments Per Kilobase of transcript per Million mapped reads (FPKM) method.

### PDAC scRNA-seq data processing

Single-cell RNA sequencing (scRNA-seq) profiles of 24,478 cells from 24 pancreatic ductal adenocarcinoma (PDAC) tumor samples were obtained from the previous study^26^. This dataset was utilized to investigate the expression levels of CpG-associated genes across distinct cell types within the tumor microenvironment. The processed gene expression matrix was imported into Seurat^48^ (v5.1.0) for downstream analysis. Stringent quality control measures were implemented to filter out low-quality cells. Cells were excluded if they met any of the following criteria: detection of fewer than 200 genes, detection of expression in fewer than three cells, or exhibition of a mitochondrial gene content exceeding 10%. After quality control filtering, the remaining high-quality cells were merged for unsupervised clustering analysis. To visualize cell clusters in a reduced dimensional space, t-distributed stochastic neighbor embedding (t-SNE) was employed. Cell clusters were then annotated and identified based on the expression of established marker genes reported in the original study.

## Author Contributions

Y.Z. and Y.Z. performed the computational analysis and data visualization. B.A.Z. conceived and designed the study, developed the methodology, wrote the manuscript.

## Competing interests

The authors have declared no competing interests.

### Acknowledgements

The National Institutes of Health supported this work: R35GM142917 and U24HG012070. Funding for open access charge: National Institutes of Health.

## Data availability

DNA methylation data were obtained from the TCGA data portal (https://portal.gdc.cancer.gov), ICGC data portal(https://dcc.icgc.org), and GEO(https://www.ncbi.nlm.nih.gov/geo/).

## Code availability

All codes used in this study are available in supplementary file 1.

