## Supplemental figures for "Machine Learning-Based Identification of Survival-Associated CpG Biomarkers in Pancreatic Ductal Adenocarcinoma"

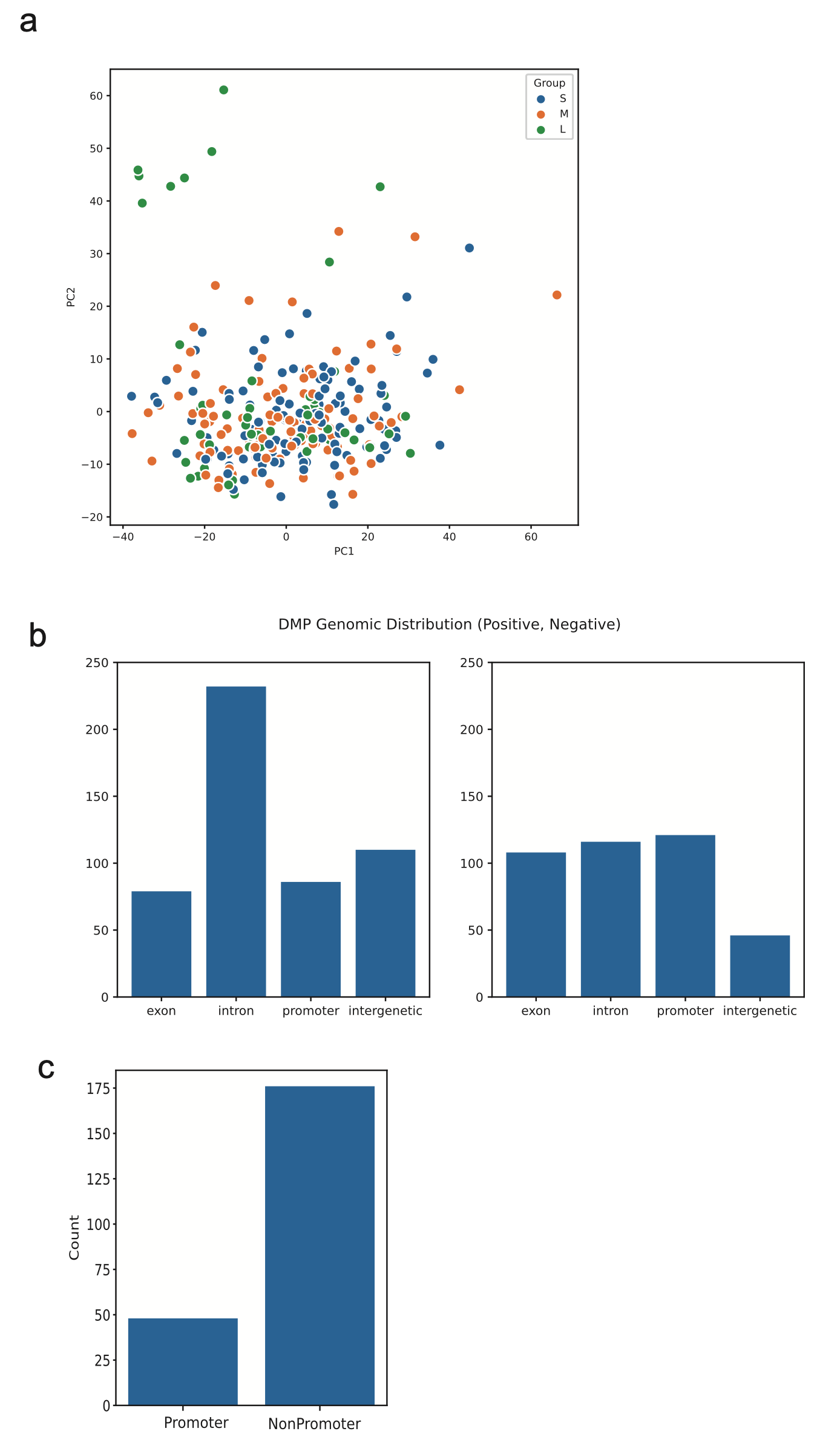

**Supplementary figure 1. a).** PCA distribution of TCGA and ICGA samples, colored by short (S), middle (M), and long (L) survival groups. **b).** Genomics distribution of 668 DMPs in human genome. Left: hypermethylated DMPs in LS group. Right: hypermethylated DMPs in SS group. **c).** number of 224 survival-associated CpG biomarkers located in the gene promoter regions.

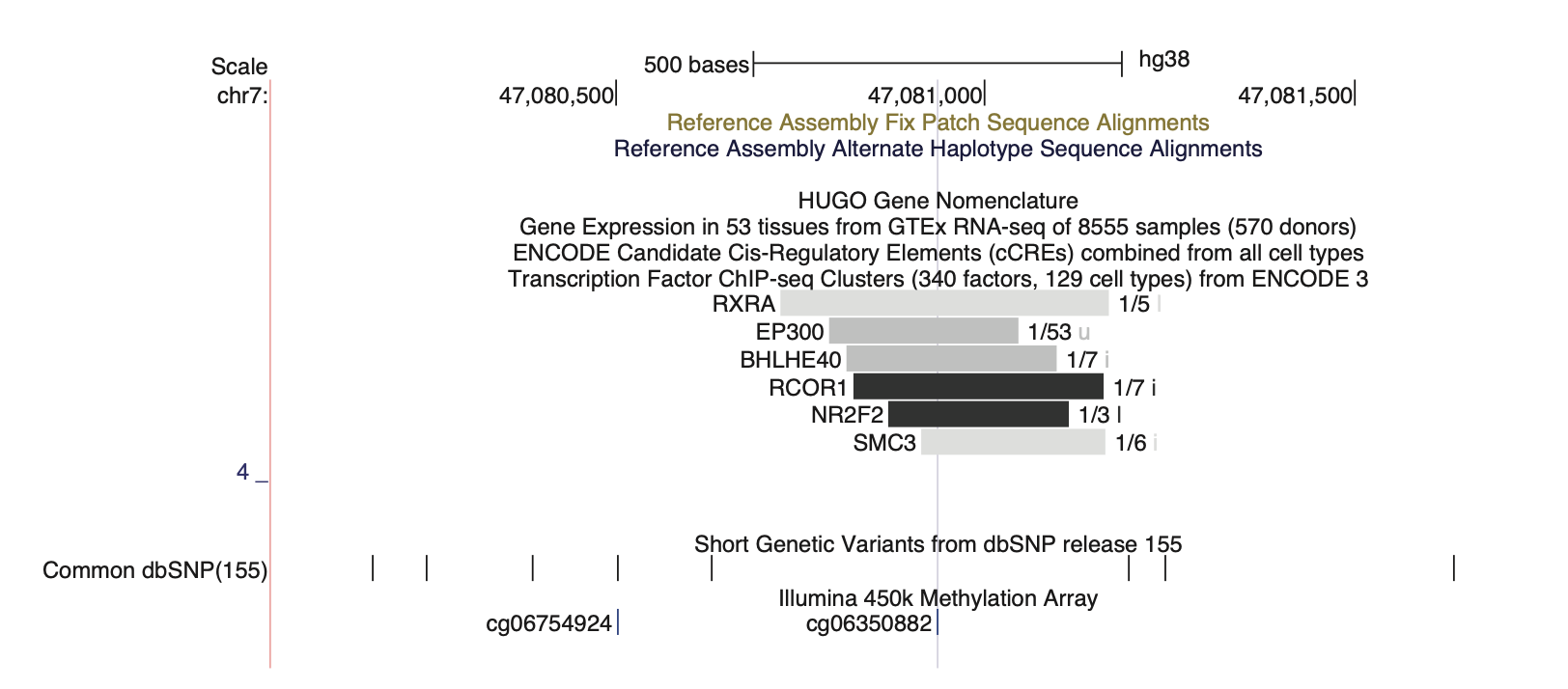

**Supplementary figure 2.** Genome browser view of cis-regulatory element Tagged by cg06350882.

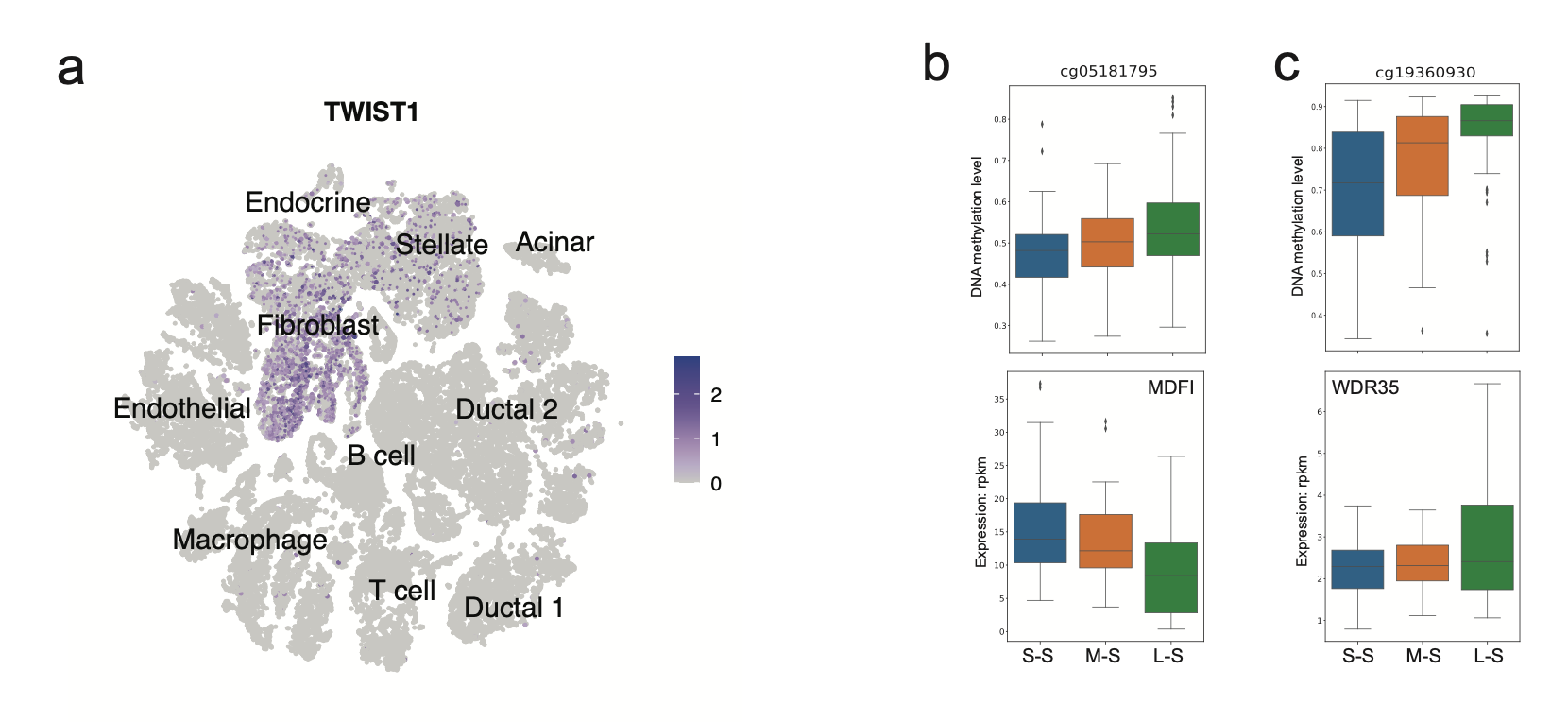

**Supplementary figure 3. a).** TWIST1 expression in distinct cell types of 24 PDAC tumors. b). DNA methylation level of cg05181795 (top) and expression level of tagged gene *MDFI* (bottom) in different patient-groups of distinct survival times. c). DNA methylation level of cg19360930 (top) and expression level of tagged gene *WDR35* (bottom) in different patient-groups of distinct survival times.

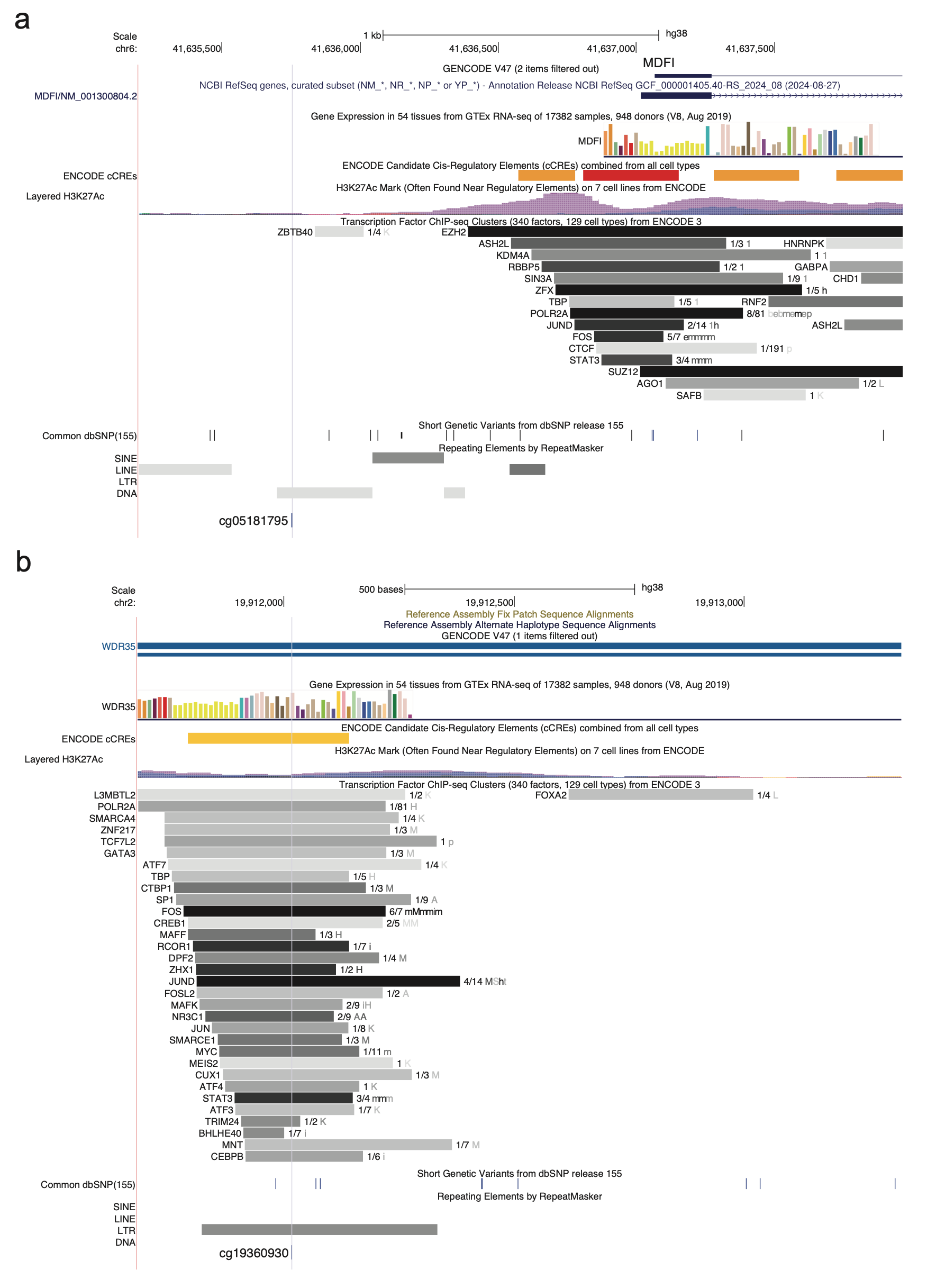

**Supplementary figure 4. a).** Genome browser view of transposable element MER96B tagged by cg05181795. **b)**. Genome browser view of transposable element LTR102_Mam tagged by cg19360930.
